# RNMT is recruited to RNA by interaction with RNA G-quadraplexes

**DOI:** 10.1101/2025.11.16.688685

**Authors:** Lydia A. Hepburn, Sander Granneman, Joana C Silva, Victoria H. Cowling

## Abstract

The 7-methylguanosine (^m7^G) cap protects RNA pol II transcripts from exonucleases and allows interaction with cap binding proteins which direct processing and translation. During^m7^G cap formation in mammals, nascent transcripts receive a guanosine cap which is methylated by RNA guanine-7 methyltransferase (RNMT). Unlike other capping enzymes which interact directly with RNA pol II, RNMT is recruited to the guanosine cap by interactions with RNA and the cap itself. RNMT is regulated during cell differentiation with significant impact on which RNAs are expressed and translated. The gene-specificity of RNMT was unexplained since RNMT in complex with activating subunit, RAM, binds to RNA without sequence preference. Through the development of improved RNA-protein detection, CLIP-ART, we report that RNMT interacts directly with RNA G4 quadraplexes (rG4). RNMT predominantly interacts with rG4s in the 5’ UTR (untranslated regions) of mRNAs, indicating a mechanism for anchoring RNMT adjacent to the guanosine cap substrate. RNMT interacts with rG4s in transcripts which encode proteins involved in growth and proliferation. The RNMT-rG4 interaction is specific to the RNMT monomer, rather than RNMT-RAM, indicating a mechanism by which differential regulation of RNMT and RAM, can lead to cap methylation of specific RNAs involved in growth control.

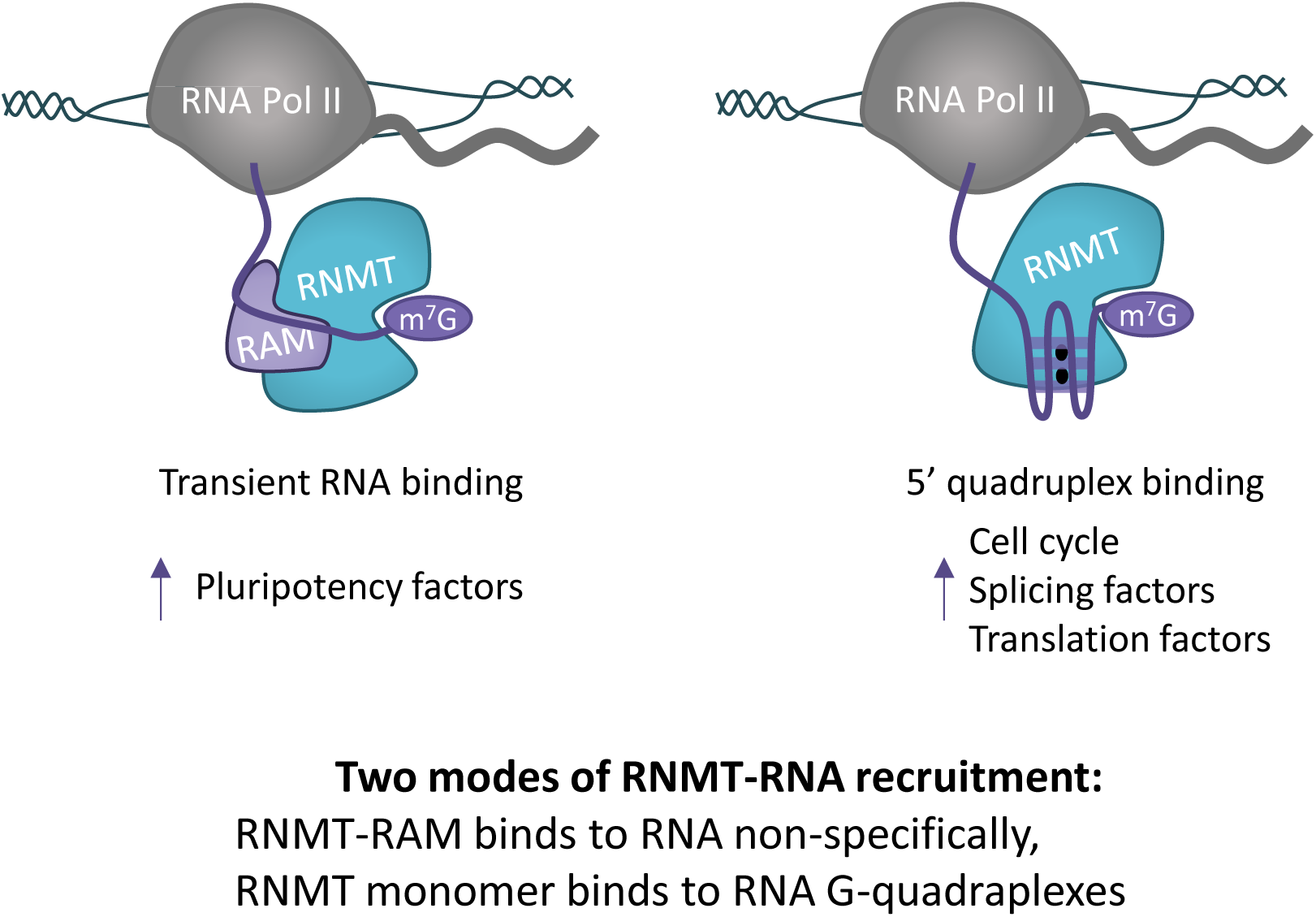
Graphical Abstract.

## Introduction

In eukaryotes, the RNA cap structure is added to RNA pol II transcripts to protect them from exonucleases and recruit factors required for processing, export and translation initiation (Galloway and Cowling, 2019, Pelletier et al., 2021). The RNA cap common to eukaryotes is 7-methylguanosine linked to the first transcribed nucleotide via a 5’ to 5’ triphosphate bridge, ^m7^GpppX (X is the first transcribed nucleotide). Although other RNA cap modifications are present, the 7-methylguanosine (^m7^G) modification has the most significant impact on RNA levels and protein production.

The RNA cap is added co-transcriptionally (Pelletier et al., 2021). An inverted guanosine group is added to the first transcribed nucleotide as it emerges from the polymerase complex, catalysed by triphosphatase and guanylyl transferase activities. In mammals, these activities are found on a single polypeptide, RNGTT (RNA guanylyl transferase and triphosphatase), which binds directly to the RNA pol II complex (Galloway and Cowling, 2019, Pelletier et al., 2021). The N-7 guanosine cap methyltransferase, RNMT, completes the ^m7^G RNA cap, ^m7^GpppX (Mills et al., 2025). A robust interaction between RNMT and RNA pol II complex proteins has not been observed; RNMT does not have a WW domain through which other RNA cap methyltransferases interact with RNA pol II. RNMT is likely to be recruited directly to the guanosine cap and via interaction with RNA (Varshney et al., 2018).

A key role of RNMT and the ^m7^G cap is to protect RNA during transcription. Deletion of RNMT results a significant reduction in mRNA per cell (Galloway et al., 2021). However, targeting of RNMT does not impact all RNAs equivalently; certain RNAs exhibit enhanced dependency on RNMT (Galloway et al., 2021, Grasso et al., 2016, Fustin et al., 2013, Del Valle Morales et al., 2020, Varshney et al., 2018). The mRNAs most dependent on RNMT vary with cell lineage, due in part to which RNAs are expressed. The level of RNMT post-translational modifications and co-factor abundance also impact on which RNAs interact with RNMT and are stabilised by receiving a ^m7^G cap. Furthermore RNMT (and homologues) have methylation-independent functions in transcription, including recruiting the PAF complex to nascent RNA and chromatin (Varshney et al., 2018).

RNMT binds to RNA directly and through the NR-rich domain within the RNMT co-factor RAM (Wen and Shatkin, 2000, Gonatopoulos-Pournatzis et al., 2011, Gonatopoulos-Pournatzis and Cowling, 2014). To understand the gene specificity of RNMT, RNMT-RNA interactions were analysed using an iCLIP protocol (Individual nucleotide CrossLink ImmunoPrecipitation) protocol (Varshney et al., 2018). RNA-RNMT interactions occurred across the transcript, appearing weak or transient and relatively non-specific.

Here, we modified a single end CLIP (se-CLIP) protocol to map where RNMT binds to mRNAs in embryonic stem cells (ESC) and neural stem cells (NSC). We demonstrate that RNMT interacts with RNA G4 quadraplexes (rG4), G-rich secondary structures within RNA. RNMT interacts with rG4s predominantly in the 5’ UTR of specific transcripts, including those associated with gene expression and proliferation. rG4s interact with the RNMT catalytic domain (but not the active site), indicating a mechanism by which RNMT is recruited to the guanosine cap. When RNMT levels are repressed in ESC, the expression of proteins encoded by transcripts with RNMT-rG4 interactions are also reduced.

## Results

### RNMT binds to secondary structures in the 5’ UTR

To map where RNMT binds to cellular transcripts, we used a modified seCLIP protocol to capture RNMT-RNA complexes in murine embryonic stem cells (ESC) and neural stem cells (NSC). The use of ESC and NSCs derived from ESC (Pollard et al., 2006), allows detection of RNMT-RNA interactions and how they change during differentiation. Cells were UV-irradiated to crosslink RNA-protein complexes, RNMT-RNA complexes were immunoprecipitated (IP) and an infrared (IR)-detected adaptor was ligated onto the 3’ ends of co-purifying RNA (**FigureS1A**) (Van Nostrand et al., 2017). RNMT and interacting RNAs were detected using western blots and infra-red, respectively (**Figure S1B**). Using this RNMT-RNA capture protocol, cDNA library synthesis with standard reverse transcription buffers was inefficient and RNA sequencing analysis revealed few RNMT binding regions (**Figure S1C and D**). RNMT-RNA binding regions that were detected were guanosine rich **(Figure S1E**). LiCl has been previously utilized in cDNA library preparations to reduce RNA G-quadruplexes, secondary structures co-ordinated by potassium or sodium ions which impeded reverse transcriptase (Kwok et al., 2016). By performing seCLIP with an Alternative Reverse Transcription (ART) buffer containing Lithium Chloride (LiCl), in a technique named seCLIP-ART, we generated complex cDNA libraries (**Figure S1A and S1C**). This indicated that a significant proportion of the RNA interacting with RNMT has secondary structure.

Inspecting crosslinks in RNMT seCLIP-ART libraries using Piranha peak calling (Uren et al., 2012) and further filtering using htseq-clips Sliding Windows (Sahadevan et al., 2022), we found that RNMT binds within 1,481 transcripts in ESC and 1,379 transcripts in NSC, with a 588 (∼40%) transcript overlap (**Figure 1A**). Pathway analysis of the RNMT-RNA interactome shows enrichment for transcripts encoding proteins involved in M-phase, pre-mRNA processing and mRNA splicing in both stem cell types (**Figure 1B**). Within identified transcripts, RNMT binding regions locate primarily within the 5’UTR and introns (**Figure 1C-E**). The use of CAGE data indicates that the peak of RNMT interaction with RNA is within 100 bases of the transcription start site (TSS) (Takahashi et al., 2012). RNMT recruitment to specific transcripts via secondary structure in the 5’ UTR may enhance interaction with its substrate, the guanosine cap. Motif analysis of RNMT-RNA binding sites reveals an enrichment of G-rich sequences in ESCs and NSCs, indicating the potential presence of RNA G-quadraplexes (rG4) (**Figure 1F**).

**Figure 1.**
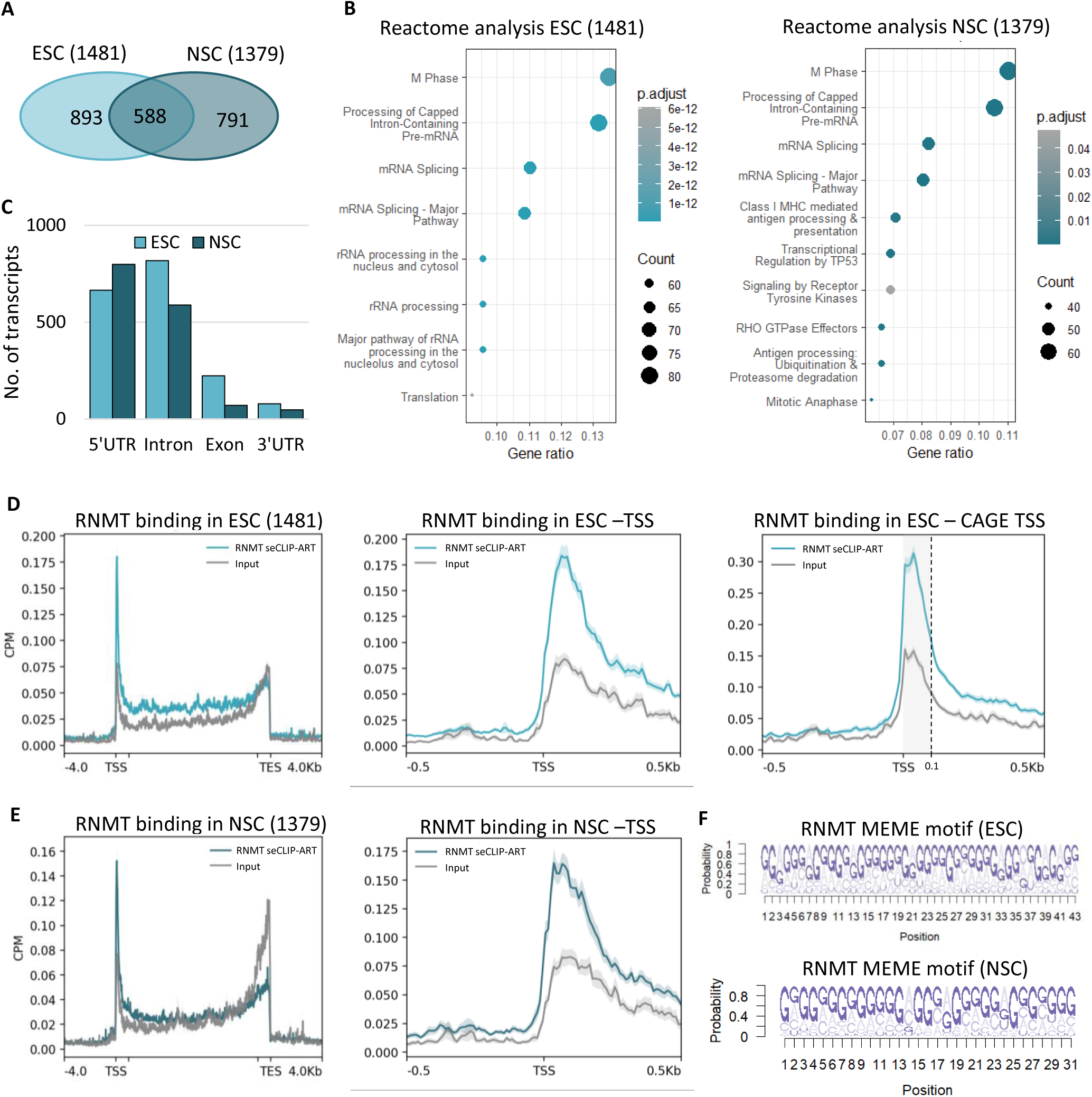
RNMT binds to G-rich motifs at the 5’ end of transcripts in ESCs and NSCs. **A**) Number of transcripts containing RNMT binding regions in ESC and NSC. Number of RNAs specific for each cell line and overlap shown. **B**) Reactome analysis of RNMT bound transcripts in ESCs and NSCs. Groups presented. Dot sizes correspond to number of transcripts in each group. **C**) Analysis of RNMT binding regions within the transcript body. **D, E**) Gene profiles for RNMT-RNA crosslinks along the gene body and 500nt around the TSS. Whole gene body is unscaled for 1kb at the TSS and TES as indicated by the x-axis notches. Blue, RNMT crosslink CPM. Grey, input CPM. D) Created using 1481 RNMT-interacting transcripts identified in ESC. E) Created using 1379 RNMT-interacting transcripts identified in NSC. **F)** MEME suit analysis of RNMT binding motifs in ESC and NSCs.

### RNMT binds to RNA G-quadruplexes in ESC and NSC

The canonical G4 consists of four repeats of three sequential guanosines, with variable loop regions of 1-7 nucleotides between each G-repeat that can form in DNA and RNA (**Figure 2A**) (Lane et al., 2008). In G4s, guanosine tetrads form into planar configurations with Hogssteen base parings, with the three planar G-tetrads pi stacking ontop of each other. Cations between the planes stabilise the structure. Additional non-canonical rG4s can form; longer loop rG4s with up to 12 nucleotides between sequential guanosines, bulge rG4s where a nucleotide breaks up the three sequential guanosines, and small G2 rG4s with four sets of two guanosines (Kwok et al., 2016). RNMT binding regions were analysed for the presence of all four types of rG4 which were defined using the method of Varshney et al, (Varshney et al., 2021). Strikingly, we found that 90% of RNMT-interacting RNA in ESC and 85% of RNMT-interacting in NSC contain one or more canonical and non-canonical rG4s within RNMT binding windows (**Figure 2B**).

**Figure 2.**
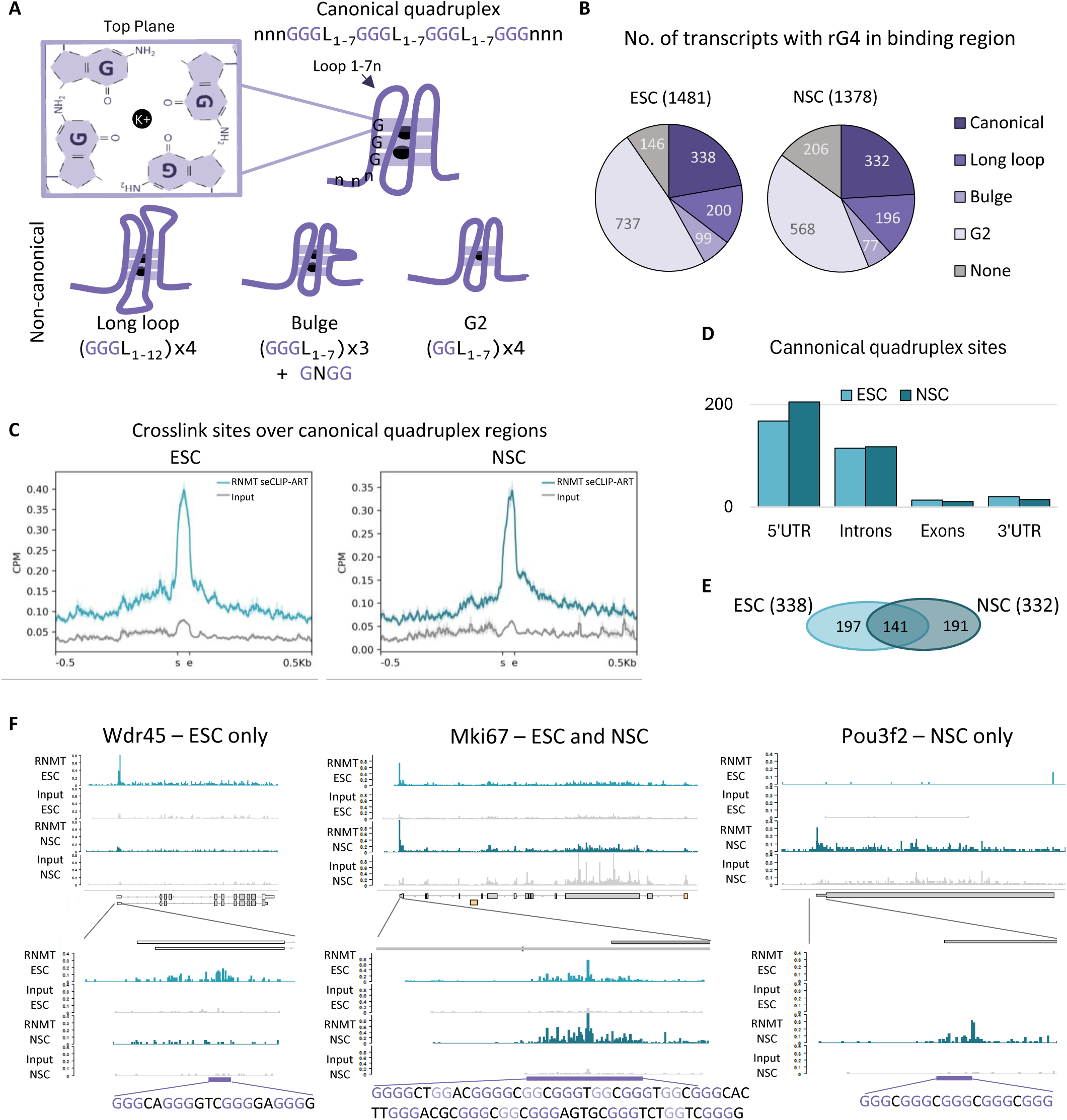
RNMT binds to RNA G-quadruplex (rG4) regions in RNAs. **A**) Canonical and non-canonical quadruplexes form from G-rich sequences. Dark purple, RNA sequence. Black, Potassium/Sodium ion. Light purple, guanine tetrad plane. **B**) Number of RNMT-bound transcripts having potential canonical and non-canonical rG4s within binding regions. **C**) Alignment of RNMT-RNA cross-links identified in seCLIP-ART, aligned with canonical rG4 sequence, in ESCs and NSCs. rG4 start(s) and end (e) depicted. Blue, seCLIP-ART CPM. Grey, input CPM. **D)** Distribution of RNMT cross-linking regions containing canonical rG4s. **E**) Number of RNMT cross-linking region with canonical rG4s in ESC and NSC. Overlap depicted. F) Individual transcripts: Wrd45, mki67 and Pou3f2. RNMT crosslink sites in CPM for ESC (light blue) and NSC (dark blue). ESC input (light grey) and NSC Input (Dark grey) plotted. Canonical rG4 (purple box) and RNA sequences with guanine highlighted in purple.

Plotting RNMT crosslink sites within 0.5kb of these RNA G-quadraplexes resulted in a peak of binding directly over the rG4 sequence (**Figure 2C**).

In ESC and NSC, RNMT binding to canonical rG4s occurs primarily within 5’UTR and introns (**Figure 2D, E, Table S1 and S2**). Detection of RNMT-rG4 interactions may partially depend on the abundance of the transcript. For example, in ESC and NSC, RNMT is found on the canonical rG4 in the 5’UTR of the mki67 transcript, a marker of proliferation (**Figure 2F**). A transcript with a RNMT-bound canonical rG4 exclusively in ESC is Wdr45, a ribosome biogenesis factor. RNMT crosslinks are found across the Wdr45 transcript, with an enrichment at the 5’UTR and within an rG4 region (**Figure 2F**). An NSC-specific transcript, Pou3f2, has RNMT bound in the 5’UTR within a canonical rG4 region (**Figure 2F**).

Of note, in the initial, low-resolution RNMT seCLIP analysis (using standard buffers during reverse transcription), crosslinks occurred across transcripts, but with no defined 5’ binding compared to the seCLIP-ART analysis (**Figure S2A and B**). RNMT binding directly over rG4s was also observed in this RNMT seCLIP, albeit with lower enrichment than in the RNMT seCLIP-ART (**Figure S2C**). RNMT seCLIP crosslinks skew to one side of the G-quadraplex consistent with premature stopping of the reverse transcriptase at secondary structure.

### RNMT interacts directly with rG4 structures

To confirm the RNMT-rG4 interaction in an orthogonal assay, we applied an affinity enrichment (AE) assay using biotinylated RNAs to purify proteins from ESC lysates. We tested two potential rG4 sequences found within the RNMT seCLIP and seCLIP-ART data sets (eEIF4a1-rG4, Snrnp70-rG4) (**Figure 3A and S2D**). For positive and negative controls, we used the characterised rG4 found within the 5’UTR of NRAS RNA (NRAS-rG4) and a mutated version (NRAS-mut) (Herdy et al., 2018). These RNA interaction assays were performed on uncapped transcripts, since RNMT-RNA interactions occur independently of the cap (Gonatopoulos-Pournatzis et al., 2011). Prior to the interaction assays, bait RNAs were resolved on TBE/acrylamide gels and stained with the rG4-specific fluorescent stain, *N*-methylmesoporphyrin IX (NMM). This confirmed formation of rG4 in EIF4a1-rG4, Snrnp70-rG4, NRAS-rG4s and a lack of rG4 in NRAS-mut RNA (**Figure S3A**).

**Figure 3.**
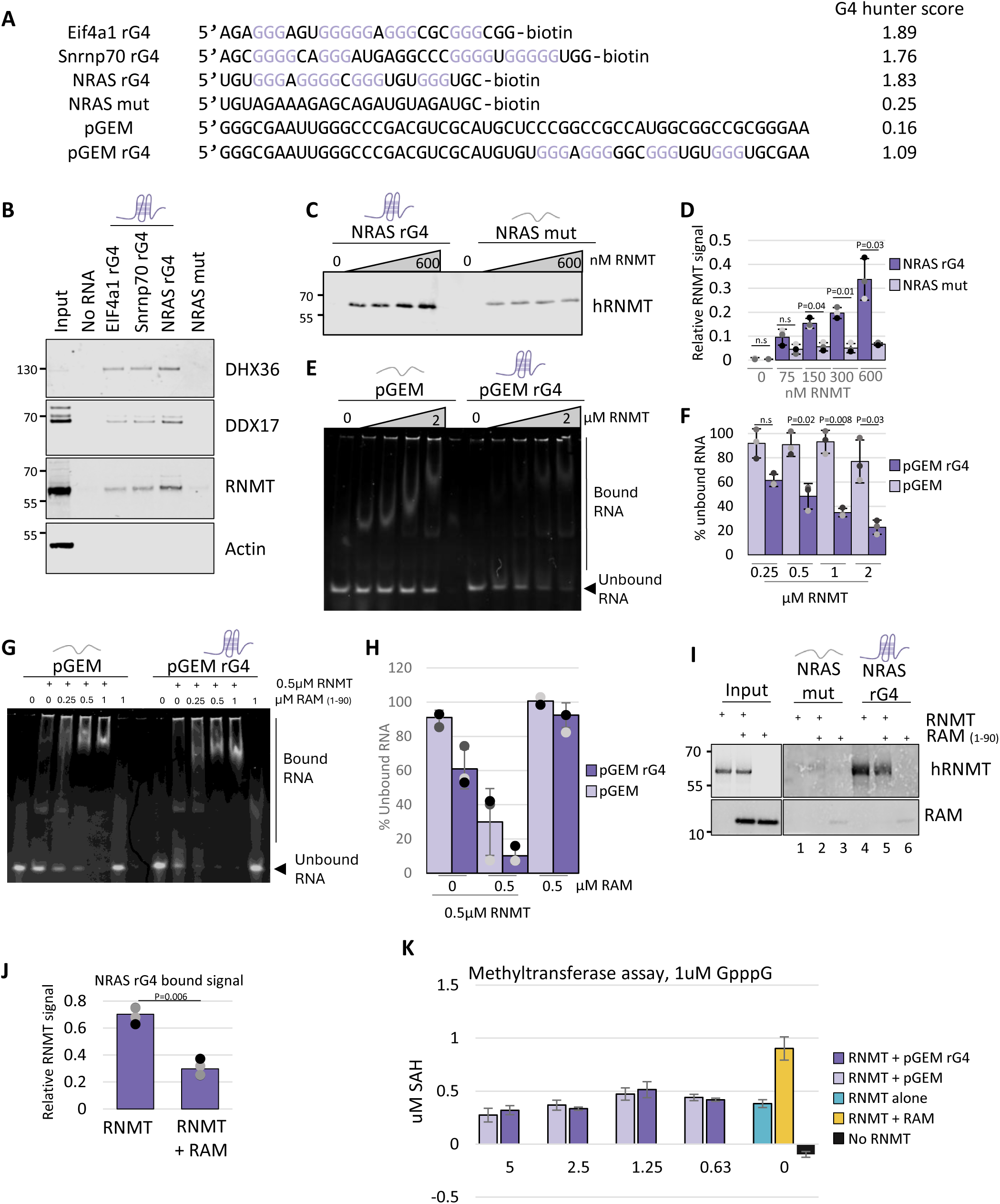
RNMT interacts with rG4 directly. **A)** RNA sequences used for affinity enrichment (AE) and EMSA, with G4 hunter score. **B**) Western blot detection of proteins binding to RNAs in AE assay. EIF4a1 rG4, Snrnp70 rG4, and NRAS rG4 contain rG4s. Actin was used as negative control. **C**) Western blot detection of human RNMT (hRNMT) protein pull downed with NRAS rG4 and NRAS mut in AE. 0, 75, 150, 300, 600 nM RNMT used. **D**) Quantification of RNMT pulled down (n=3). Dark purple, NRAS rG4. Light purple, NRAS mut. **E)** Representative EMSA of pGEM RNA or pGEM rG4 RNA incubated with a titration of recombinant h RNMT (0, 0.25, 0.5, 1, 2 µM RNMT). Gel stained with SYBR gold. Arrow indicates unbound RNA fraction. **F)** Percentage of unbound RNA in EMSA (n=3). Dark purple, pGEM rG4 RNA. Light purple, pGEM RNA. **G)** Representative EMSA with pGEM RNA or pGEM rG4 RNAs and recombinant human RNMT and RAM (1-90aa). Gel stained with SYBR gold. Arrow indicates unbound RNA fraction. **H)** Percentage of unbound RNA in EMSA. Dark purple, pGEM-rG4 RNA. Light purple, pGEM RNA(n=3). **I)** AE of NRAS RNA and NRAS-mut RNA with recombinant RNMT, +/− RAM (1-90aa). **J)** Quantification of NRAS-RNMT interaction +/− RAM (n=4) **I)**) **K**) Methyltransferase assay with stated concentration of RNA. 500nM RAM added to the reaction of 500nM RNMT, 1µM GpppG and 2µM SAM.

In rG4 AE assays, ESC lysates and biotinylated RNAs were incubated together, then RNA-protein complexes were washed and proteins eluted. Established NRAS-rG4 interactors, DHX36 and DDX17, bound to the NRAS-rG4 and both novel RNMT-interacting rG4s, EIF4a1-rG4 and Snrnp70-rG4 (**Figure 3B**). RNMT pulled down with all rG4s, confirming the specificity of RNMT-rG4 interactions identified by RNMT seCLIP-ART. We also tested binding of recombinant human RNMT (hRNMT) to the NRAS-rG4 RNA. hRNMT interacts with NRAS-rG4 RNA in a concentration-dependent manner, whereas binding to NRAS-mut RNA was minimal (**Figure 3C, D**). This indicates that RNMT binds directly to rG4 structures, independently of auxiliary proteins. RNMT-RNA interactions have been analysed by RNA electromobility band shift assay (EMSA) (Gonatopoulos-Pournatzis et al., 2011, Wen and Shatkin, 2000). We performed an EMSA with recombinant hRNMT and *in vitro* transcribed RNA from the pGEM-T plasmid, either the pGEM sequence or an rG4-containing transcript, pGEM-rG4. The presence of rG4 in the pGEM-rG4 transcript was confirmed by NMM staining (**Figure S3A**). In the EMSA, detection of RNMT-RNA complexes as band shifts was inconsistent between replicates (**Figure 3E, S3D**).

However, over a titration of RNMT, the amount of unbound, free rG4-RNA reduced, consistent with RNMT-rG4 interaction and the RNMT-RNA complexes smearing throughout the gel (**Figure 3E, F**). The unbound control-RNA was only reduced at the highest concentration of RNMT, consistent with reduced RNMT binding is the absence of rG4.

### RNMT-rG4 interaction is inhibited by co-activator RAM

Previously, we observed that the RNMT co-activator RAM binds to RNA in an EMSA and enhances non-specific RNA recruitment to RNMT (Gonatopoulos-Pournatzis et al., 2011). Here, we also observe that RNMT and RNMT-RAM bind to RNA in an EMSA (**Figure 3G, H**). RNMT-RAM binds non-specifically to the pGEM RNA with or without the rG4 (Gonatopoulos-Pournatzis et al., 2011, Varshney et al., 2018). (RAM 1-90, rather than full length RAM 1-118 is used throughout, because full-length RAM 1-118 is unstable (Gonatopoulos-Pournatzis et al., 2011)).

EMSAs are performed with low salt and detergent, thus capture low affinity protein-RNA interactions. The AE is a more stringent protein-RNA interaction assay compared to the EMSA, including detergent wash steps, thus weaker interactions are lost. Under the stringent conditions of this assay, when present alone RAM exhibits minimal interaction with NRAS-rG4 or NRAS-mut RNA (**Figure 3I**, lanes 3 and 6; RAM detection). Of note, the presence of RAM reduces RNMT interaction with NRAS-rG4 (Figure 3I,compare lanes 4 and 5, RNMT detection). Even when present with RNMT, RAM binding to NRAS-rG4 was not detected above background (NRAS mut) **(Figure 3I lane 4, RAM detection**), indicating that RNMT monomer and not RNMT-RAM is binding to rG4. Therefore, although RAM increases RNMT cap methyltransferase activity (Gonatopoulos-Pournatzis et al., 2011) and non-specific RNA binding (**Figure 3G and H**), it reduces RNMT interaction with rG4 (**Figure 3I and J**).

To investigate whether RNMT-rG4 interactions alter RNMT cap methyltransferase activity an *in vitro* assay was performed. Of note, in this assay, the rG4 RNA is not capped: this assay is not measuring whether the presence of the rG4 increases methylation of the cap on the same RNA. RNMT was incubated with substrate GpppG, methyl donor SAM (s-adenosyl methionine) and a titration of rG4-RNA and control-RNA (both uncapped). Methylation was detected by the production of SAH (S-adenosyl homocysteine) in the absence of RNA (**Figure 3K)**. The presence of rG4 or control RNA did not alter the catalytic activity of RNMT. Furthermore, internal ^m7^G methylation of these G-rich was not detected and was not expected due to the shape of the RNMT active site (Varshney et al., 2016). RAM, the RNMT activator was included as a positive control in these activity assays.

### rG4 interaction occurs within the catalytic domain of RNMT

Human RNMT has a C-terminal catalytic domain (120-476) and a N-terminal regulatory domain (1-120) which impacts on catalytic activity and recruitment to chromatin (Aregger and Cowling, 2013, Aregger et al., 2016) (**Figure 4A**). The catalytic domain contains the active site and RAM binding site (Varshney et al., 2016). To investigate which parts of RNMT are utilised in RNMT-rG4 interactions, we generated four recombinant RNMT fragments, amino acids 1-152, 1-185, 120-476 and 165-476 (**Figure 4A and B**). RNMT fragments were used in RNA-interaction assays (AE and EMSA). In AE, RNMT 1-185 and 120-476 bind to rG4-RNA with enhanced affinity compared with bindingcontrol-RNA (**Figure 4C, S4A and B**). Similarly, in the RNMT EMSA, RNMT 1-185 binds to rG4-RNA with higher affinity than control-RNA (loss of free rG4 probe). RNMT 120-476 binds to both rG4-RNA and control-RNA (**Figure 4D, E and S4B**). RNMT 1-152 and 165-476 do not exhibit enhanced binding to rG4 RNA. Taken together, these data indicates that the rG4 is likely to interact with or be dependent on RNMT 152-165.

**Figure 4.**
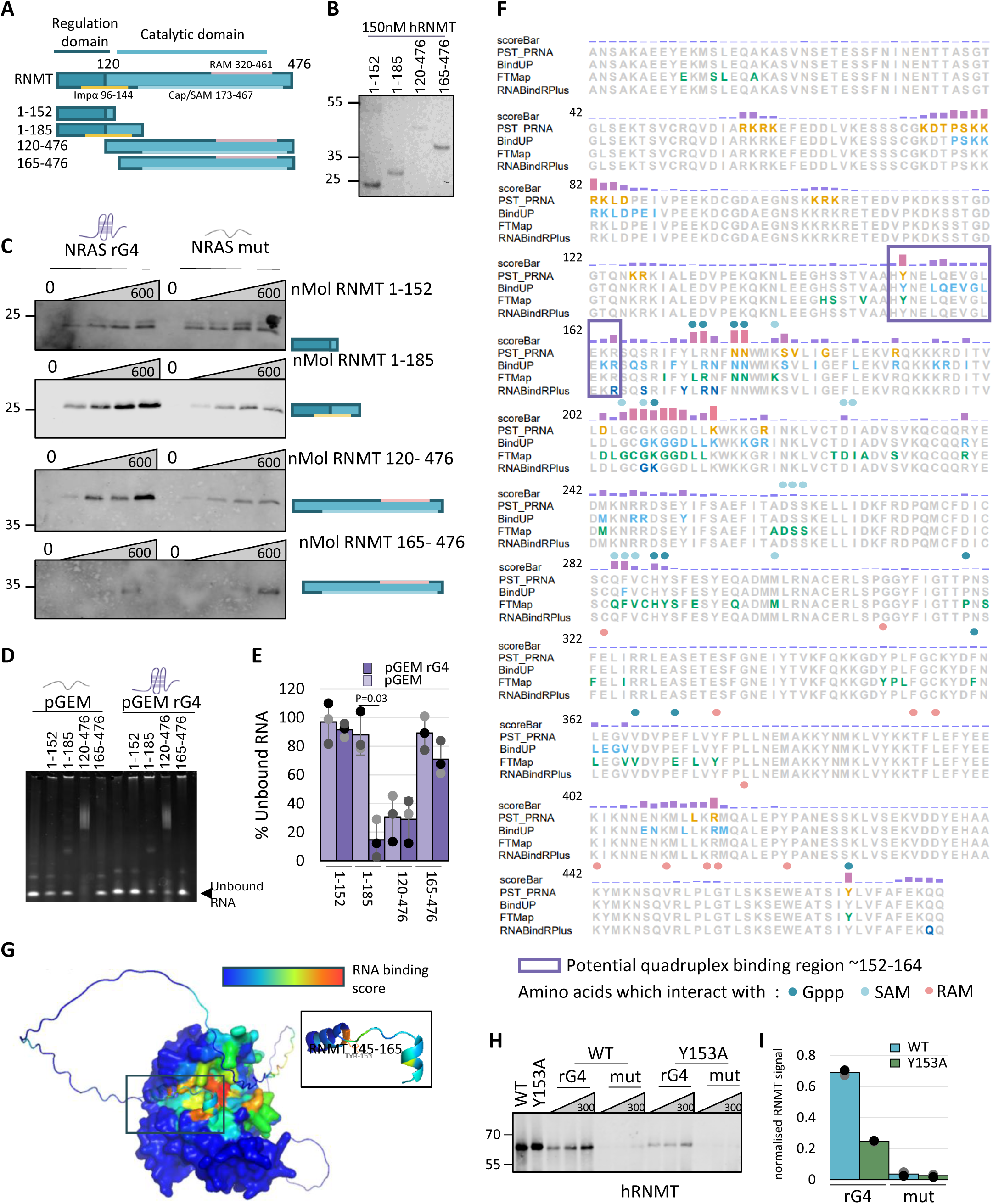
RNMT catalytic domain binds to rG4. **A)** Human RNMT domains. hRNMT trunctions utilised. **B)** Coomassie Blue-stained gel of recombinant hRNMT truncations. **C)** Recombinant hRNMT truncation proteins pulled down with NRAS rG4 RNA and NRAS mut RNA, in AE. **D**) Representative EMSA with pGEM RNA and pGEM rG4 RNA and hRNMT truncations. SYBR gold-stained gel. **E**) Percentage of unbound RNA (n=3). Dark purple, pGEM rG4 RNA. Light purple, pGEM RNA. **F)** pyRBDome results for human RNMT. Score bar indicates likelihood of RNA interactions. Amino acids predicted to interact with the GpppG cap cap (Dark blue), SAM (light blue) and RAM (purple) highlighted. Potential rG4-interacting region, purple box. **G)** AlphaFold predicted structure for hRNMT with pyRBDome RNA binding score indicated. Dark blue (low) to red (high). **H**) AE pulldowns with NRAS rG4 RNA or NRAS mut RNA and 75, 150 and 300 nM hRNMT WT or Y153A. RNMT Western Blot. **I**) Quantitation of RNMT pulldown at 300nM RNMT. Blue, WT. Green, Y153A mutant (n=2).

The human RNMT amino acid sequence was analysed pyRBDome, a Python-based computational pipeline that uses published crystal structures, Alphafold predictions and multiple distinct computational tools to predict RNA-binding residues based on electrostatic potential, sequence conservation and protein structure (Jumper et al., 2021). An XGBoost ensemble model is then employed to create a final score that indicates for each amino acid residue a probability that it binds RNA (Chu et al., 2024) (**Figure 4F**). Amino acids which interact with the RNA cap and SAM (methyl donor) within the active site have a high pyRBDome RNA-binding probability, consistent with nucleotide binding. Within the region mapped as a potential rG4 interaction site (**Figure 4A-D**), there are multiple amino acids which score highly for sites of potential RNA interaction, including Y153, Q157 and R164 (Figure 4E). This region of RNMT has not been crystallised, but Alphafold predicts the formation of two small alpha helices and places the region over the catalytic domain (**Figure 4G**). Since Y153 has the highest pyRBDome score, we generated the RNMT Y153A mutant. Consistent with a deficiency for RNA binding, RNMT Y153A binds with reduced affinity to NRAS-rG4 RNA compared to the control (**Figure 4H and I**).

### Targeting of RNMT results in repression of rG4 containing proteins

rG4s have been demonstrated to have multiple impacts on RNA dynamics (Dumas et al., 2021). In ESC, we targeted RNMT expression to investigate the impact of RNMT-rG4s interactions. To focus on the role of RNMT-rG4 interactions, ESC in which part of the RAM gene is deleted using CRISPR were used for this analysis (RAM^Del^, **Figure S5A**). In RAM^Del^ ESC, there is a 21 nucleotide in-frame deletion within the coding region of RAM **(Figure S5a**). RAM protein is not detected in RAM^Del^ ESC extracts or RNMT IPs and there is decreased RNMT, consistent with RAM stabilising RNMT protein (**Figure S5B-D**)(Gonatopoulos-Pournatzis and Cowling, 2014). As observed previously, when RAM levels are reduced in ESC, Oct4 expression is reduced whereas Nanog is maintained (**Figure S5B**) (Grasso et al., 2016). Although the amount of RNMT is reduced in RAM^Del^ ESC, when levels of RNMT are balanced, the amount of RNA crosslinked to RNMT is does not decrease (RNA analysis of RNMT CLIP-ART samples, **Figure S5D**).

In RAM^Del^ ESC we investigated multiple RNMT-bound transcripts. Smc1 was used as a control since RNMT crosslinks are enriched within exons but not at the 5’ UTR (Figure 5A, B).

**Figure 5.**
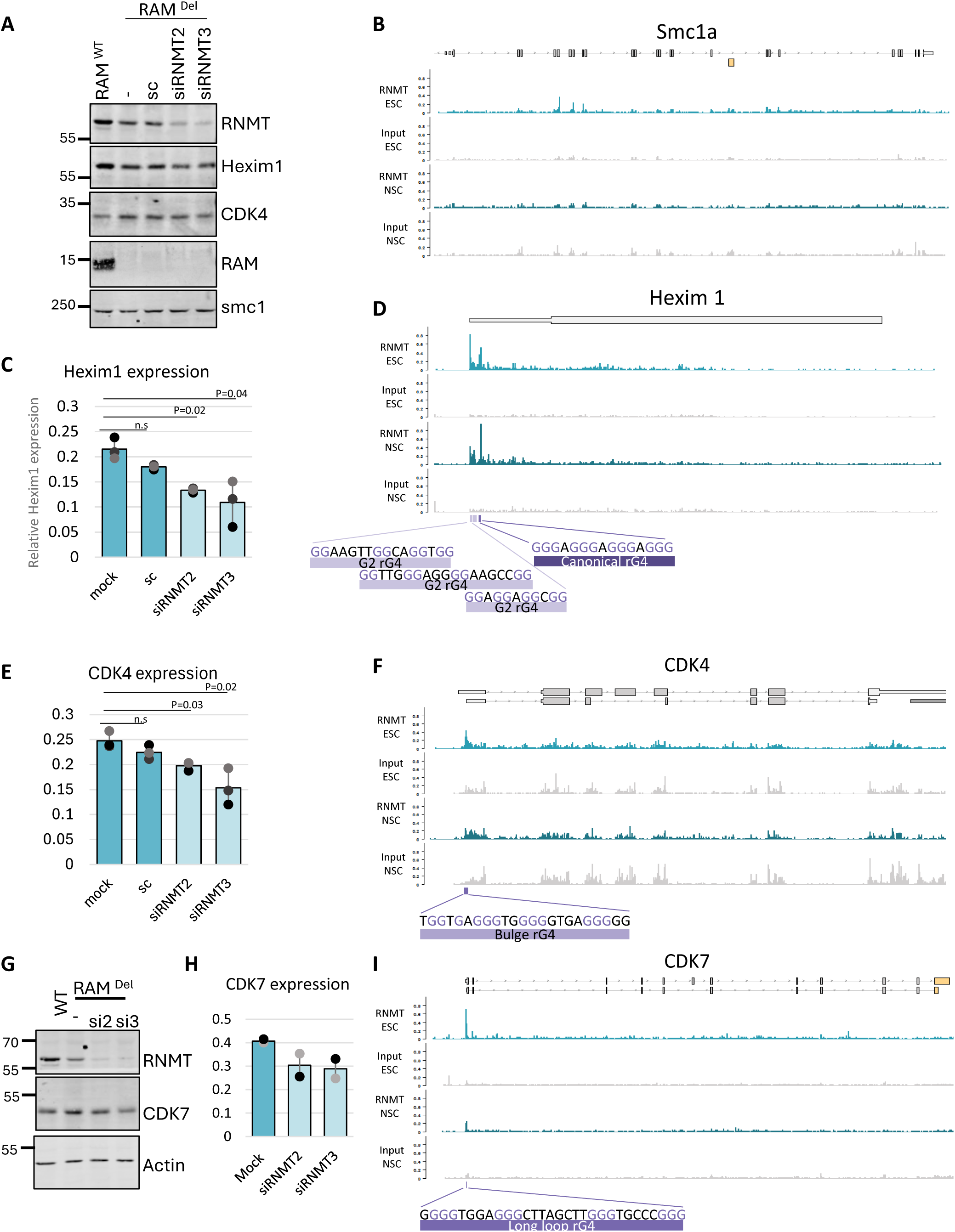
**For mRNA with RNMT-rG4 interaction, RNMT knockdown reduces protein product**. Representative western blot for indicated proteins in RAM ^WT^ or RAM ^Del^ cells with siRNMT B) RNMT crosslink sites from seCLIP-ART in ESCs and NSCs for smc1a, y=CPM. C) Quantitation of hexim1 expression western blots (n=3). D) RNMT seCLIP-ART crosslink sites across Hexim1, 5’ UTR rG4 sites highlighted in purple, y=CPM. E) Quantitation of CDK4 expression western blots. F) Crosslinks or RNMT seCLIP-ART across CDK4 with highlighted rG4 sequence, y=CPM. G) Representative western blot of siRNMT knockdown and indicated proteins. H) Quantitation of CDK7 expression western blots. I) Crosslinks of RNMT seCLIP-ART across CDK7, Quadruplex site highlighted, y= CPM

Knockdown of RNMT had no detectable effect on SMC1 protein (**Figure 5A and B**). RNMT crosslinks to the 5’UTR of Hexim1, which contains, three G2 rG4s and a canonical rG4. When RNMT is reduced in the RAM^Del^ ESC using RNMT siRNA, Hexim1 protein is reduced (**Figure 5C and D**). RNMT binds to the CDK4 mRNA bulge rG4 within the 5’UTR and to CDK7 long loop rG4 (Figure 5E and F). RNMT targeting also results in reduction of CDK7 and CDK4 protein (**Figure 5 E and F**).

## Discussion

Here we report that the ^m7^G RNA cap methyltransferase, RNMT, interacts directly with rG4s. RNMT binds to rG4s predominantly in the 5’UTR, proximal to the initial transcribed nucleotides and the unmethylated guanosine cap. Since RNMT binds to rG4 in introns, this indicates at least some binding during transcription. RNMT has a co-factor RAM which increases catalytic activity; RAM inhibits RNMT binding to rG4s; under stringent conditions rG4s interact with RNMT and not with RAM or RNMT-RAM. This indicates that RNMT-rG4 interactions are specific to RNMT monomer, which was previously thought to be simply a RNMT protein with reduced activity. We now recognise that RNMT monomer is likely to have unique target RNAs and functions, independent of RNMT-RAM.

### Exploring the CLIP-ART technique

The observation of rG4s interact with RNMT became possible due to adaptations in the seCLIP reverse transcriptase step. RNA secondary structures stall and impede reverse transcriptase and although increased temperature reduces secondary structures, rG4s can stably reform in the presence of KCl or NaCl (Hardin et al., 1992). Following the observation of poor library formation and G-rich sequences in the RNMT seCLIP, the reaction buffer salt was exchanged to LiCl, which had been previously utilised in other CLIP-related methods to permit reverse transcription through rG4 structures (Varshney et al., 2021). This CLIP-ART protocol (with LiCl-based buffers) resulted in the mapping of RNMT-rG4 interactions in ESC and NSC.

### Exploration of the RNMT rG4 interaction

During screening projects to identify rG4 binders, RNMT was identified in rG4– and dG4-interactomes, using several cell lines and using different bait G-quadruplex sequences. RNMT was found in telomeric dG4-interactomes in Hela cells (Pipier et al., 2021), NRAS rG4-interactomes in Hela cells (Herdy et al., 2018), and in rG4-interactomes in U251 glioblastoma cells (Herviou et al., 2020). We demonstrate a direction interaction between RNMT and rG4, but these prior studies suggest that RNMT can interact with a broad range of quadruplex sequences, beyond rG4 and across cell lineages.

The region of RNMT which interacts with rG4 maps to the N-terminus of the catalytic domain, a region with little structural characterisation. Alphafold analysis indicates that this RNMT region forms an alpha helix, and a single tyrosine mutation within this region, Y153A, is disrupts rG4 binding (Jumper et al., 2021). The G-quadruplex helicase, DHX36, contains the DHX36-specific motif (DSM), a 13 amino acid alpha helix containing a tyrosine which pi-stacks on top of the exposed guanosine tetrad plane (Banco et al., 2025, Chen et al., 2018). Mutation of this tyrosine is sufficient to dissociate the DSM from the quadruplex (Chen et al., 2018). The DHX36 DSM is intrinsically disordered and only forms an alpha helix upon interaction with a G-quadruplex (Heddi et al., 2015). Potentially co-crystalising RNMT with an rG4 may allow for the structure of the interaction region to be stabilised.

### rG4s in the transcriptome

The mapping of rG4s within the transcriptome has been investigated using rG4-seq, RT-stop profiling, and a modified SHAPE-seq (Kwok et al., 2016, Guo and Bartel, 2016, Zhao et al., 2025). These rG4 mapping techniques utilise reverse transcriptase stalling in KCl to map which G-rich sequences will form the secondary structure. These studies reveal rG4s to form primarily within the 3’UTR, however this may be partially influenced by the 3’ bias of poly A tail selection of the RNAs profiled (Cui et al., 2010). UV-RNA Immunoprecipitation using a rG4-specific antibody, found greater enrichment of rG4 within the 5’UTR than the 3’ UTR or coding sequences (Varshney et al., 2021).

rG4 sequences within 5’UTRs are enriched around sites of RNA pol II pausing (Eddy et al., 2011, Rosenberg et al., 2021). ESCs exhibit high levels of RNA pol II pausing, particularly on cell cycle-related genes; mutations which inhibit pausing reduce differentiation (Tastemel et al., 2017, Williams et al., 2015). Here we demonstrate a propensity for RNMT to bind to these RNA pol II-enriched cell cycle transcripts in ESC and NSCs. In Hela cells, targeting of RNMT in Hela cells reduces RNA pol II occupancy, particularly around the 5’ TSS, where pausing occurs (Varshney et al., 2018).

In T cells, *Rnmt* deletion reduces the expression level TOP RNAs, which contain a terminal oligopyrimide (TOP) motif in their 5′UTR (Galloway et al., 2021). TOP RNAs are in part regulated by the TOP and cap binding protein, LARP1. LARP1 also interacts with rG4s and many TOP RNAs have rG4s adjacent to their TOP motif (Herdy et al., 2018, Herviou et al., 2020, Pipier et al., 2021, Varshney et al., 2021). RNMT may also be being recruited these 5’ rG4s in TOP-RNAs and influence cap methylation or other aspects of RNA dynamics.

### rG4-interacting methyltransferases

Multiple RNA methyltransferases have also been found to interact with rG4s, with a range of impacts. m6A methyltransferase, METTL3/14, interacts with rG4s through the RGG domain of METTL14, which directs methylation to adenosines found within loop regions or close by rG4 sequences (Yoshida et al., 2022). The m7G methyltransferase, METTL1, methylates rG4 sequences within precursor miRNA, which counteracts the inhibitory activity of rG4 on miRNA processing. BUB23, TRMT112 and NSUN5 which methylate rRNA, were also identified in rG4 interactomes (Herdy et al., 2018, Herviou et al., 2020). It is not known which of these enzymes interact directly with rG4s, or the impact of rG4 binding.

### Control of RNM-rG4 interactions by RAM

We do not detect rG4 interaction with the RNMT activator, RAM, or the RNMT-RAM complex in affinity enrichment assays *in vitro.* RNMT and RAM expression vary in different tissues and RAM is targeted for degradation during neural differentiation (Grasso et al., 2016). RAM is also upregulated during T cell activation and during the later stages of neural differentiation (Galloway et al., 2021). The ratio of RNMT monomer to RNMT-RAM is likely to influence RNMT-rG4 interactions in these scenarios, with impacts depending on the positioning of the rG4.

### Summary

In summary, we report that RNMT interacts directly with rG4s. rG4s map to a region within the catalytic domain of RNMT. RNMT rG4 interaction occurs predominantly within the 5’UTR of transcripts, near the cap, potentially increasing cap methylation or another co-transcriptional mechanism. rG4s in introns, coding regions and 3’UTRs may have other impacts on RNA dynamics. The RNMT-rG4 interaction indicates a method of recruitment of to specific transcripts and may be the key to understanding the gene specificity of RNMT in different cellular contexts.

## Methods

### Cell culture

Embryonic Stem cells (ESC) containing SOX1 promoter-driven GFP from strain 129Ola mice were grown on 0.1% gelatin-coated plates in Glasgow MEM (ThermoFisher Scientific) containing 10% Knockout serum replacement (ThermoFisher Scientific), 1X Non-essential amino acids (NEAA) (ThermoFisher Scientific), 2mM L-glutamine (ThermoFisher Scientific), 0.1mM β-mecaptoethanol and recombinant human Leukemia Inhibitory Factor (LIF) 100U/ml (DSTT).

Neural Stem cells (NSC) were created from the SOX1:GFP cells, as previously described (Pollard et al., 2006) and cultivated on Matrigel-coated plates in DMEM/F12 (ThermoFisher Scientific) containing with 1.4mg/ml Glucose (Sigma), 2mM L-glutamine, 1X NEAA, 0.5X B27 (ThermoFisher Scientific), 0.5X N2 (ThermoFisher Scientific) and 0.1mM β-mecaptoethanol and supplemented with fresh 100µg/ml Fibroblast Growth Factor-basic (Peprotech) and 100µg/ml murine Epidermal Growth Factor (Peprotech).

### SiRNA transfection

Cells (7.5X10^5^) were reverse transfected with 120pMol of siRNA (Darmacon) using 2.4µl lipofectamine RNAiMAX (ThermoFisher Scientific) reagent. Collected 48hrs after transfection

### CRISPR/Cas9 genome targeting

Antisense gRNA against RAMAC, cloned into pX335-Cas9-D10A. Sense gRNA cloned into pBABED-Puro-U6. ESC cells at ∼70% confluency were reverse transfected using Lipofectamine 2000 (Life Technologies) and 1 μg of each gRNAs vector. Cells were selected using 1 μg/ml puromycin for 2 days, then left to recover. Low density allowed single colonies to form. Single colonies were then selected and analysed for protein expression and DNA sequenced.

### seCLIP-ART analysis

Library preparation: Cells were washed with ice cold PBS before being UV crosslinked (UVitech) at 400 mJ/cm^2^. Cells were lysed in CLIP lysis buffer (Tris HCl pH 7.5 50mM, NaCl 100mM, SDS 0.1%, NP-40, 1%, Sodium deoxycholate 0.5%, supplemented with 5.5µl/ml protease inhibitor III (NEB), 11µl/ml RNAse murine inhibitor (NEB) and 20µl/ml TURBO DNase (ThermoFisher)), The lysate was diluted to 1µg/µl and 2 mg of protein per IP digested with Ambion RNAse I (100U/mL) diluted to 1:20,000 and incubated for 5 mins at 37°C. Lysates were left mixing overnight at 4°C with 6µg of α-RNMT sheep antibody (DSTT) and Protein G Dynabeads (ThermoFisher Scientific). 2% inputs were boiled in LDS (ThermoFisher scientific), 100mM DTT and samples processed as previously described (Blue et al., 2022) with the following exceptions. For RNA imaging, the DNA adaptor /5Phos/CAAGCAGAAGACGGCATACGAAAAAAAAAAAA/iAzideN/AAAAAAAAAAAA labelled with DBCO 800 dye (licor) was ligated to a portion of the IP RNA-RNMT complexes with T4 RNA ligase 1 (NEB). All IPs and inputs were resolved on 4-12% NuPAGE™ Bis-Tris Protein Gels (ThermoFisher Scientific) in 1X MOPS buffer (ThermoFisher Scientific) and transferred onto nitrocellulose membrane (Amersham). The membrane was imaged on the Licor Clx in the 800 channel. Western blot was performed with the α-RNMT rabbit primary Ab Antibody (DSTT) and 680 anti-rabbit (Licor) secondary Ab. Immediately after transfer of the sequencing IP portion, the membrane was cut between 55 to 130 kDa and protease-digested (as previously) to release the RNA from the protein-RNA complexes. RNA was isolated using RNA Clean and Concentrator-5 (Zymo Research) and then reverse transcription with SuperScript III (ThermoFisher Scientific) was carried out in the Superscript Buffer as seCLIP or in the 5X ART buffer (100mM Tris HCl pH 7.5, 16mM MgCl_2_, 750mM LiCl) (Kwok et al., 2016, Varshney et al., 2021).

#### Read processing

Libraries were run on Novaseq with 150pb single-end sequencing. Reads containing unique molecular identifiers (UMIs) were extracted from the sequencing data using Umi tools extract. Adaptors were removed from reads using cutadapt. Reads were aligned to the Mm10 version of the mouse genome with version M25 gencode gene annotation using STAR align. Aligned reads were sorted and indexed using samtools. Duplicate reads were removed using umicollapse tool (Liu, 2019). The deduplicated reads were then sorted.

#### Binding analysis

Crosslink enrichment was carried out using the Piranha tool (Uren et al., 2012). Specifically, Piranha was set up using a bin size of 200, 0 cluster distance and a p value of 0.0001. Each IP was individually analysed using the extracted crosslink site bed file, windows were only taken forward which were found in all IP outputs. Crosslink numbers for each window were processed using subread and windows which contained an average of 1CPM and 2-fold CPM IP over input reads were kept. To create more refined windows of crosslinks for better annotation, Htseq-clip (Sahadevan et al., 2022) was used to create a sliding window reference file of the mm10 genome into windows of 100nt with a slide of 20 nt. The crosslink sites were extracted from the processed reads using htseq-clip extract with the first nucleotide and stranded, creating a bed file. Counts of the crosslink sites per window were created using htseq-clip count generating a matrix of the number of crosslink sites that are found within each window of the reference. Crosslink counts per million (CPM) of each IP and input sample were then averaged (average of four samples for CLIP, and three samples for CLIP-ART) and any windows which did not have at least 1CPM and averaged 2-fold CPM over input were removed. The sliding windows were then merged if they overlapped. Merged sliding Windows which were found to overlap with the Pirhana windows were then taken forward for annotation. Any windows which were not found to overlap from either method were removed. Genome enrichment analysis was carried out using the ReactomePA R package (Yu and He, 2016).

#### Gene profile

Accumulative bam files were made from all IP samples or Input bam files using samtools merge. Bigwig files were made from these bam files using bamCoverage extracting the first nucleotide as the crosslink site and normalised to counts per million. Gene profiles were made using bigwigs with deeptools computeMatrix with the scaled transcript size of 10000 nt, unscaled 1000nt at the 5’ and 3’ ends and a maximum threshold of 10 CPM then plotProfile using the mean and with standard error applied Quadruplex profiles were created using scripts as detailed in (Varshney et al., 2021). Single gene profiles were made using the GVIZ R package (Hahne and Ivanek, 2016).

### Synthetic RNAs

Biotinylated RNAs were synthesised by IDT (Integrated DNA Technologies). Other RNAs were *in vitro* transcribed using pGEM T easy vector (Promega) as a template, either with its original sequence or with a quadruplex sequence cloned into pGEM using *Apa1* and *EcoR1* restriction sites. Prior *in vitro* transcription with T7 polymerase (Promega), vectors were linearised with *EcoRI.* After IVT, RNA was purified using RNA clean and concentrator (ZymoResearch) and a micro-biospin chromatography column (Bio-Rad). To ensure preserving formation of quadruplexes, all RNAs were diluted to 1µM in 150mM KCl, denatured at 95°C for 5 mins and left to cool at 0.1°C/sec to RT before use.

### Affinity enrichment (AE)

Streptavidin Dynabeads M-280 (ThermoFisher Scientific) were washed with PBS, incubated with 5 nM biotinylated RNAs for 30 mins at RT and washed with binding buffer before use. For whole cell lysate affinity enrichment, cells were lysed in the cell binding buffer (50mM Tris pH 7.5, 100mM NaCl, 0.2% NP-40), diluted to 1µg/µl protein. 200µg lysates were incubated with 20µl of bead-RNA mixture for 2 hours at 4°C. Next, beads were washed three times with the binding buffer and eluted by boiling for 5 minutes in 2X LDS with 100mM Dithiothreitol (DTT). RNMT was detected by western blot, as previously. For recombinant protein AE, proteins were made up in a recombinant binding buffer (20mM Tris pH 7.5, 100mM NaCl, 20% glycerol, 0.2nM EDTA, 0.01% NP-40). Next, proteins were incubated with the bead-RNA mixture for 2 hours at 4°C. Then, beads were washed three times with a wash buffer (20mM Tris pH 7.5, 100mM NaCl, 20% glycerol, 0.2nM EDTA, 0.1% NP-40). Proteins were eluted by boiling at 95°C for 5 minutes in 2X LDS with 100mM DTT and analysed by western blot.

### Electrophoretic mobility shift assay (EMSA)

Recombinant proteins were diluted in EMSA buffer (20mM Tris pH 7.5, 100mM NaCl, 0.2nM EDTA, 0.01% NP-40) and incubated with RNAs for 10 mins at RT. Samples were mixed with 6X DNA Sample buffer (ThermoFisher scientific) and resolved on 4-20% TBE gels (ThermoFisher Scientific) in 1X TBE buffer at 4°C. Gels were stained with SybrGold (ThermoFisher) for 30 mins and imaged on Biorad Chemidoc.

### RNA cap methyltransferase assay

Using Promega MTase Glow assay with recombinant RNMT and 1µM GpppG synthetic cap substrate (NEB) and 2µM SAM. Protein and cap were incubated for 30mins and then heat inactivated at 70°C for 10mins. MTase glow reagent was then added to 1X and incubated for 30mins. Detection reagent added on top and luminescence reading taken after 30 minute incubation.

### Statistical analysis

Graphpad prism was used for all statistics shown and were achieved using paired Student t-tests. Image analysis including blots and gels was carried out using ImageJ software.

### Data availability

E-MTAB-16237

## Acknowledgement

We thank Dhaval Varshney, Fiona Haward, Olga Suska and past and present members of the Cowling lab. We thank Core Services and the Advanced Technologies at Cancer Research UK Scotland Institute (A31287). The research was funded by a Wellcome Trust

Investigator award 219416/Z/19/Z; a European Union Horizon 2020 European Research Council Award 769080 TCAPS. This work was supported by Cancer Research UK core grant number A17196/A31287, CRUK Scotland Institute, and CTRQQR-2021∖100006 to CRUK Scotland Centre.

## Figure Legends

**Figure S1.**
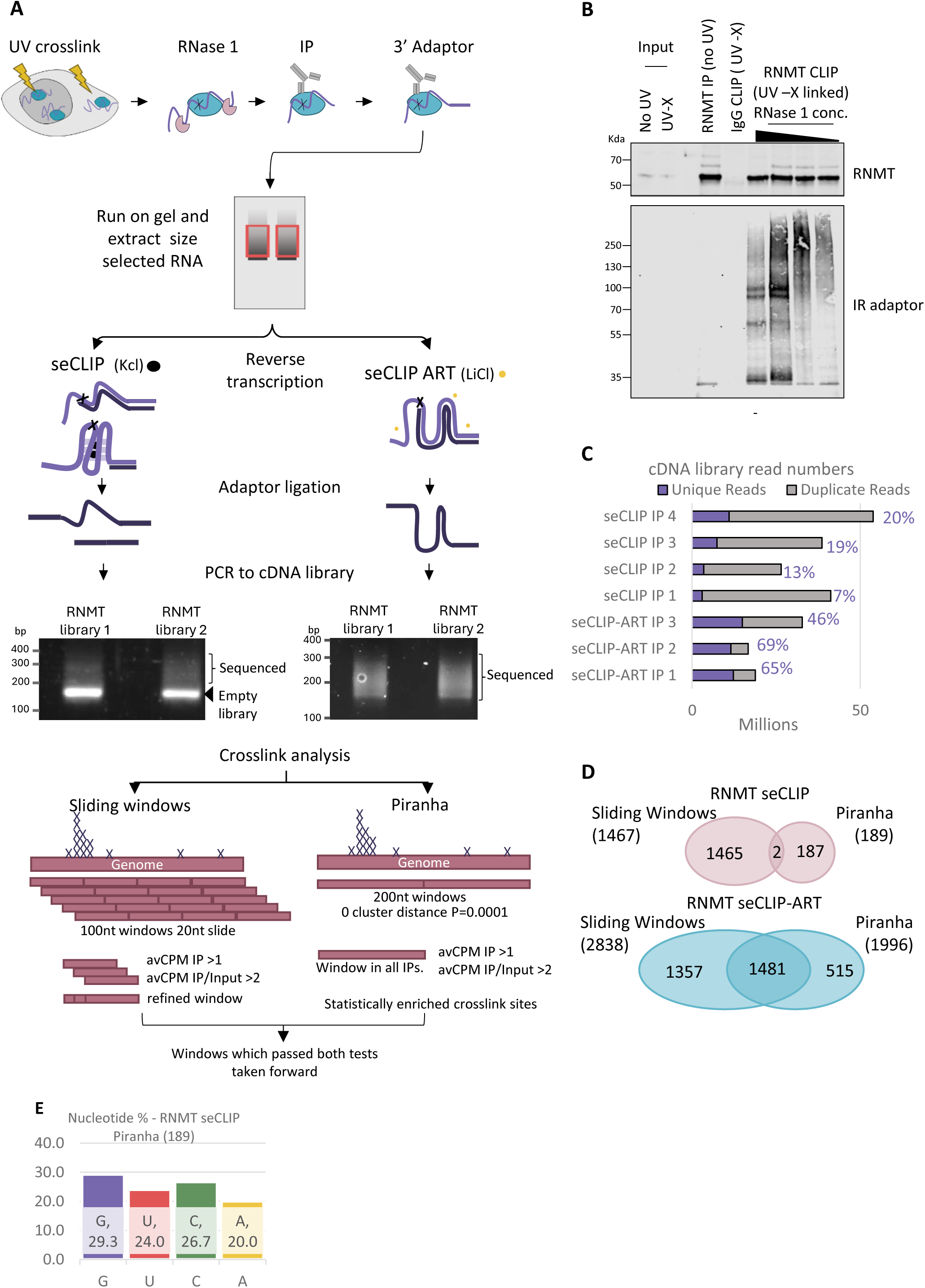
Establishment of CLIP-ART protocol: generation of complex cDNA libraries containing G-rich sequences. **A**) Overview of the seCLIP protocol. Cells are exposed to UV which crosslinks RNA (purple) to protein (blue). Cells are lysed and treated with RNase1. RNMT is immunoprecipitated (IP). The 3’ of RNMT-bound RNA is ligated to an adaptor. RNA-RNMT complexes are resolved on SDS-PAGE gel and transferred to nitrocellulose. RNA-RNMT complexes are cut out from the membrane at the region above migration of RNMT alone and RNA is extracted. In the seCLIP protocol, reverse transcription stage utilises KCl buffer. In the seCLIP-ART adaptation, an alternative reverse transcription (ART) buffer containing LiCl was used, promoting unfolding of rG4 secondary structures, which would otherwise inhibit the reverse transcriptase. The reverse transcriptase stalls at the RNMT-RNA crosslink site. An adaptor is ligated to the 3’ end of the cDNA and cDNA is amplified by PCR and analysed on gel. The cDNA library is size selected and sequenced. Crosslink analysis of unique reads was carried out by two methods: a sliding windows approach where the genome was separated into 100 nucleotide (nt) windows with 20nt between each window, and using the peak caller, Piranha, which separates the genome into 200nt windows and calls regions where there are statistically significant clusters of crosslink sites. The number of reads/crosslink sites are counted in each window and counts per million (CPM) were calculated for each sample. Only windows which have an average CPM of all IP samples over one and two-fold over input were taken as positive for RNMT binding. **B**) Visualisation of RNA crosslinked to RNMT and optimisation of RNAse1 concentration. Ligation of the adaptor containing fluorophore to 3’ end of RNA crosslinked to RNMT allows complex visualisation. An IgG antibody IP and non-crosslinked samples are used as negative controls. Input samples are 1% of the material used for IP. **C**) Percentage of reads (per million of reads) from seCLIP (four replicates) and seCLIP-ART (three replicates) sequenced libraries. Purple and number, percentage of unique reads. Grey, percentage of duplicated reads. **D)** Binding sites of RNMT in seCLIP and seCLIP-ART libraries identified using sliding windows and Piranha approaches. **E**) Nucleotide analysis of RNMT seCLIP Piranha windows.

**Figure S2.**
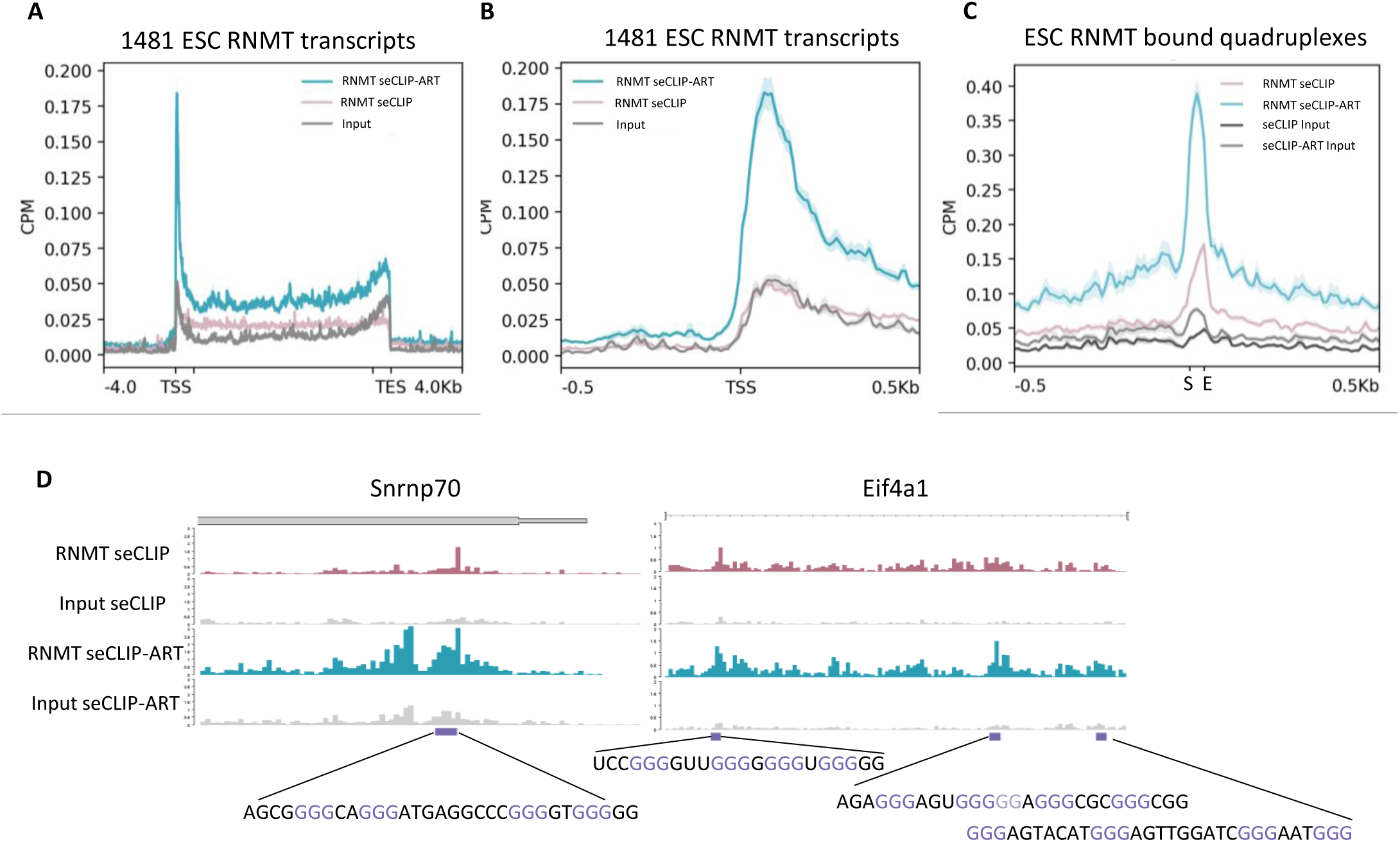
A comparison of RNMT seCLIP and RNMT CLIP-ART data. Gene profile of **A)** RNMT CLIP and input crosslinks over 1481 transcripts found bound in RNMT CLIP-ART. **B**) RNMT CLIP and RNMT CLIP-ART crosslinks over 1481 transcripts found bound in RNMT CLIP-ART. **C**) RNMT CLIP, RNMT CLIP-ART and respective input crosslinks over RNMT bound canonical quadruplex sites found with RNMT CLIP-ART. **D**) Potential rG4 sequences found within the RNMT seCLIP and seCLIP-ART data sets

**Figure S3.**
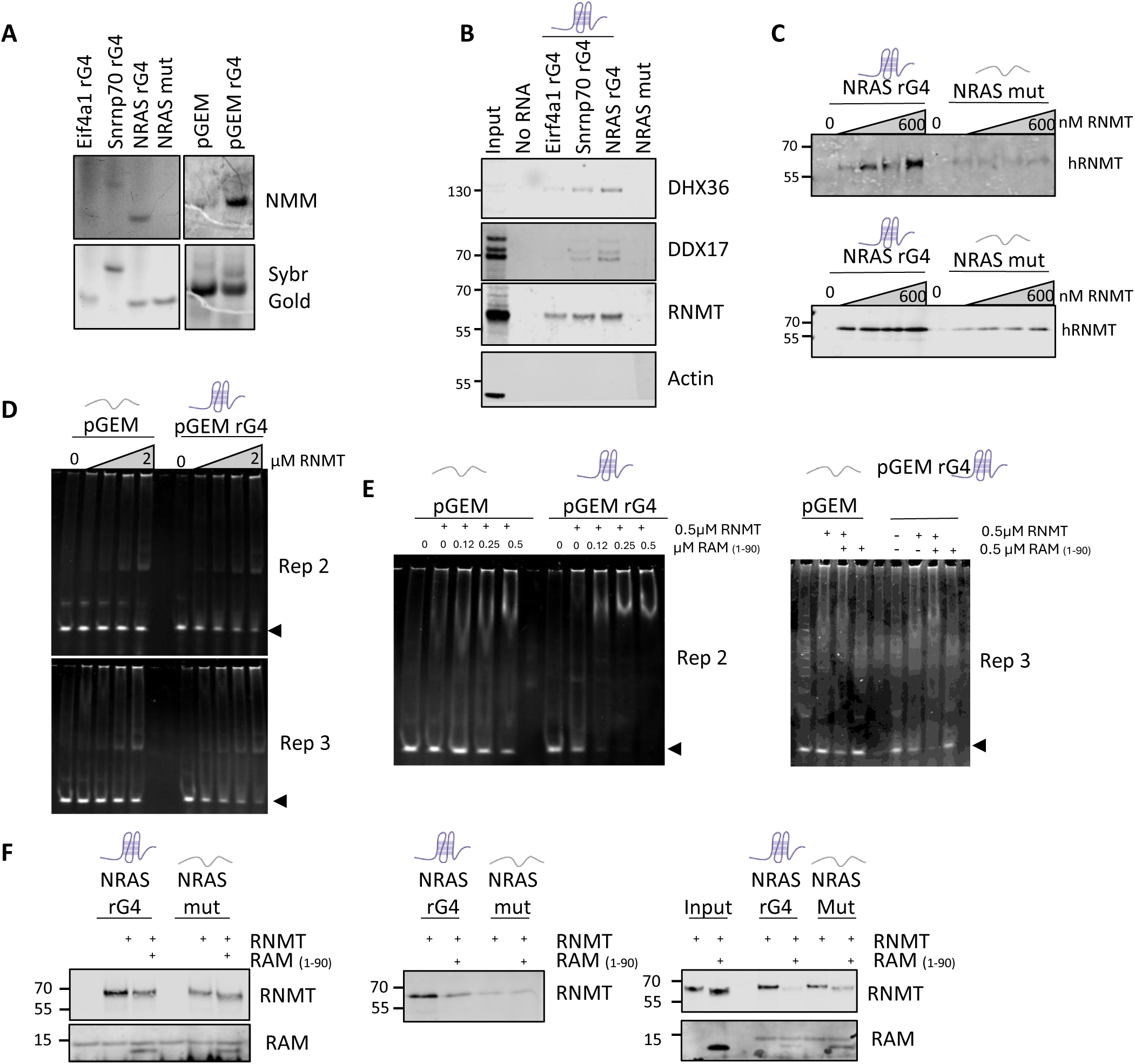
Replicate data for figure 3. **A)** NMM and SYBR gold staining of synthetic RNA resolved on gels. **B**) Replicate of RNA AE enrichment from ESC extracts. **C)** Replicates of recombinant RNMT pulldowns. **D**) Replicate EMSAs, incubating recombinant RNMT with pGEM RNA or pGEM rG4 RNA. Arrow indicates unbound RNA. **E)** EMSA repeats of RNMT and a titration of RAM1-90 with pGEM RNA or rG4 pGEM RNA. Arrow indicates unbound RNA fraction. **F)** Replicate AE pulldowns of RNMT alone or incubated with RAM.

**Figure S4.**
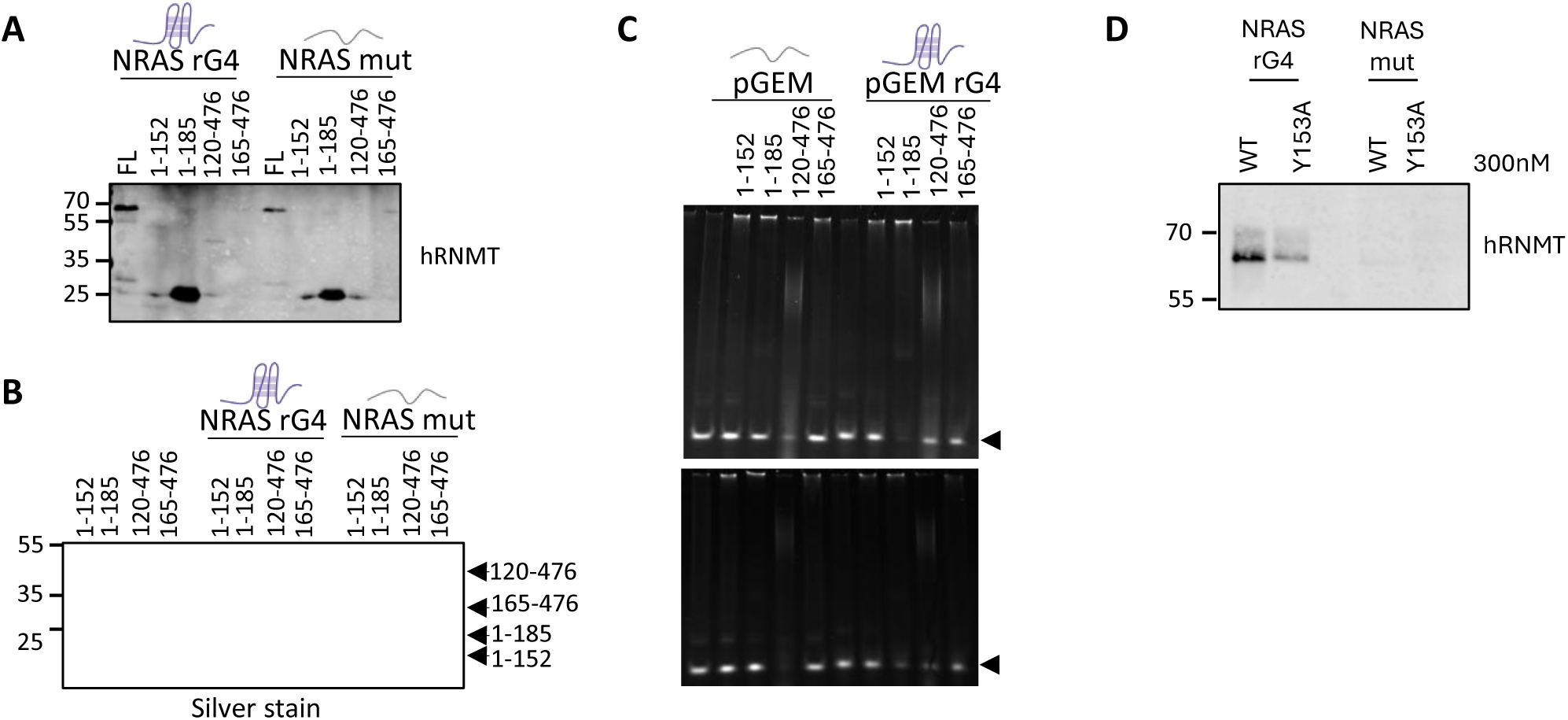
Replicate data for figure 4. **A and B**) RNMT Western blot (A) and silver stain (B) of AE pulldown with NRAS rG4 RNA or NRAS mut RNA incubated with different RNMT truncations. **C**) Replicate EMSAs of RNMT truncations incubated with pGEM RNA and pGEM rG4 RNA. **D**) Replicate western blot of AE pulldown with NRAS rG4 RNA or NRAS mut RNA incubated with recombinant RNMT WT or Y153A.

**Figure S5.**
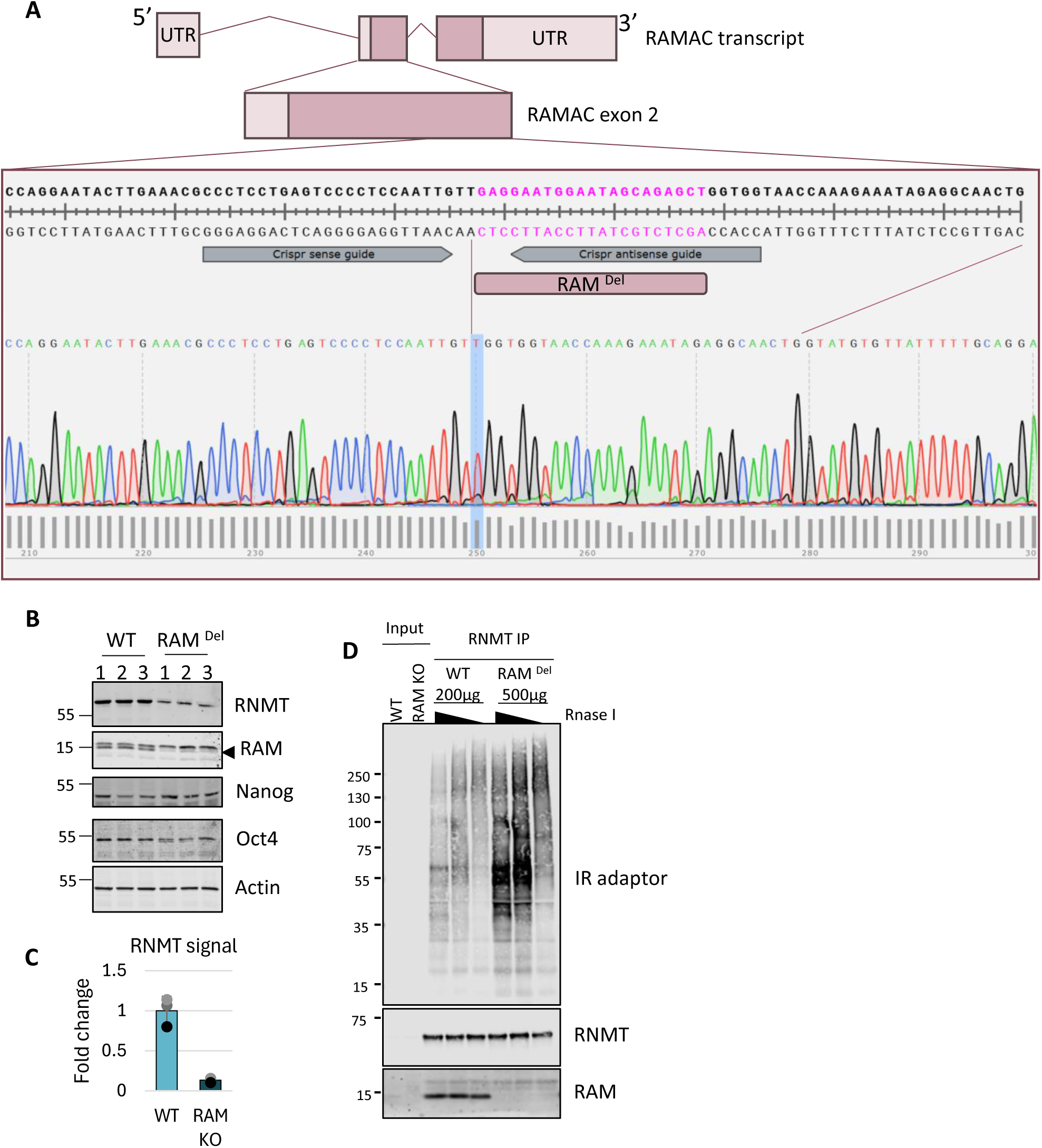
Generation of RAM^Del^ ESC. **A)** Schematic of CRIPSR guides. Sample DNA sequencing data from RAM^Del^ ESC, highlighting deleted sequence. **B**) Western blot analysis of three different passages of WT and RAM^Del^ ESCs. **C**) Quantitation of RNMT signal from B. **D**) IR CLIP of RNMT from WT and RAM^Del^ ESCs, with decreasing RNA concentration (1:2000,1:5000,1:20000 amount). RNMT IP balanced by using 200µg WT ESC extract and 500µg RAM^Del^ extract.

Table S1 ESC RNMT CLIP-ART binding windows

Table S2 NSC RNMT CLIP-ART binding windows

